# S-conLSH: Alignment-free gapped mapping of noisy long reads

**DOI:** 10.1101/801118

**Authors:** Angana Chakraborty, Burkhard Morgenstern, Sanghamitra Bandyopadhyay

## Abstract

**Motivation:** The advancement of SMRT technology has unfolded new opportunities of genome analysis with its longer read length and low GC bias. Alignment of the reads to their appropriate positions in the respective reference genome is the first but costliest step of any analysis pipeline based on SMRT sequencing. However, the state-of-the-art aligners often fail to identify distant homologies due to lack of conserved regions, caused by frequent genetic duplication and recombination. Therefore, we developed a novel alignment-free method of sequence mapping that is fast and accurate.

**Results:** We present a new mapper called S-conLSH that uses **S**paced **con**text based **L**ocality **S**ensitive **H**ashing. With multiple spaced patterns, S-conLSH facilitates a gapped mapping of noisy long reads to the corresponding target locations of a reference genome. We have examined the performance of the proposed method on 5 different real and simulated datasets. S-conLSH is at least 2 times faster than the state-of-the-art alignment-based methods. It achieves a sensitivity of 99%, without using any traditional base-to-base alignment, on human simulated sequence data. By default, S-conLSH provides an alignment-free mapping in PAF format. However, it has an option of generating aligned output as SAM-file, if it is required for any downstream processing.

**Availability:** The source code of our software is freely available at https://github.com/anganachakraborty/S-conLSH

## 1 Introduction

Single molecule real time (SMRT) sequencing developed by Pacific Biosciences [27] and Oxford nanopore technologies [21] have started to replace previous short length next generation sequencing (NGS) technologies. These new technologies have enabled us to address many unsolved problems regarding genetic variations. With the increase in read length to around 20KB [2], SMRT reads can be used to resolve ambiguities in read mapping caused by repetitive regions. Low GC bias and the ability to detect DNA methylation [27] from native DNA made SMRT data appealing for many real life applications. However, the high sequencing error rate of 13-15% per base [2] poses a real challenge in sequence analysis. Specialized methods like BWA-MEM [15], BLASR [6], rHAT [20], Minimap2 [17], lordFAST [9], etc., have been designed to align noisy long reads back to the respective reference genomes. BLASR [6] clusters the matched words from the reads and genome after indexing using suffix arrays or BWT-FM [28]. It uses a probability-based error optimization technique to find the alignment. BWA-MEM [15], originally designed for short read mapping, has been extended for PacBio and Oxford nanopore reads (with option -x pacbio and -x ont2d respectively) by efficient seeding and chaining of short exact matches. However, both methods are too slow to achieve a desired level of sensitivity [20]. This issue was addressed by rHAT [20] using a regional hash table where windows from the reference genome with the highest k-mer matches are chosen as candidate sites for further extension using a direct acyclic graph. Unfortunately, this method has a large memory footprint if used with the default word length of *k =* 13, and it fails to accommodate longer *k*-mers to resolve repeats. Minimap2 [17], a recently developed method, uses concave gap cost, efficient chaining and fast implementation using SSE or NEON instructions to align reads with high sensitivity and speed. Another new method lordFAST [9] has been introduced to align PacBio’s continuous long reads with improved accuracy. MUMmer4 [23], a versatile genome alignment system, also has an option for PacBio read alignment (-l 15 -c 31), although it is less sensitive and accurate than the specialized aligners.

However, all the above mentioned methods come with large computational costs. Here, time and memory consumption are dominated by the alignment overhead. On top of that, alignment algorithms are often unable to correctly align distant homologs in the “twilight zone” with 20-35% sequence identity, as such weak similarities are difficult to distinguish from random similarities. For these reasons, alignment-free methods have become popular in recent years. See [4, 26, 29] for recent review papers and [30] for a systematic evaluation of these approaches. An alignment-free method Minimap [16] has been developed in 2016 for mapping of reads to the appropriate positions in the reference genome. Minimap groups approximate colinear hits using single linkage clustering to map the reads. However, Minimap suffers from low specificity. In this article, a new alignment-free method called S-conLSH has been proposed to overcome the above mentioned problems. Being suitable for low conserved areas and less computationally expensive, S-conLSH is sensitive as well as very fast at the same time.

A large proportion of the sequencing errors in SMRT data are indels rather than mismatches [2]. This makes it even more complicated to differentiate genomic variations from sequence errors. To resolve this issue, a concept of ‘context-based’ Locality Sensitive Hashing (conLSH) has been introduced by Chakraborty and Bandyopadhyay [8]. Locality Sensitive Hashing (LSH)[1, 12] has been successfully applied in many real life-science applications, ranging from genome comparison [5, 7] to large scale genome assembly [3]. In LSH, points close to each other in the feature space are hashed into localized slots of the hash table. However, in practice, the neighborhood or *context* of an object plays a key role in measuring its similarity to another object. Chakraborty and Bandyopadhyay[8] have shown that contexts of symbols (a base in reference to DNA) are important to decide the closeness of strings. They proposed conLSH to group sequences in localized slots of the hash table if they share a common context. However, a match for the entire *context* is a stringent criterion, considering the error profile of SMRT data. Even a mismatch or indel of length one, caused by a sequencing error, may mislead the aligner.

Therefore, to address this problem, an idea of *spaced*-context is introduced in this article. Unlike conLSH [8] which produces base-level alignments of the sequences using Sparse Dynamic Programming (SDP) based algorithm, the proposed method (S-conLSH) is an alignment-free tool. It employs multiple spaced-seeds or patterns to find gapped mappings of noisy SMRT reads to reference genomes. The spaced-seeds are strings of 0’s and 1’s where ‘1’ represents the match position and ‘0’ denotes don’t care position where matching in the symbols is not mandatory. The substring formed by extracting the symbols corresponding to the ‘1’ positions in the pattern is defined as the *spaced-context* of a sequence. Therefore, a spaced-context can minimize the effect of erroneous bases and, thereby, enhances the quality of mapping because it does not check all the bases for a match. This differentiates the proposed method from conLSH which looks into the entire context to compute the hash values.

A pattern-based approach was originally proposed by [22] when they developed Pattern-Hunter, a fast and sensitive homology search tool. Later, multiple patterns or “spaced seeds” were proposed by the same authors [19]. Efficient algorithms to find optimal sets of patterns have been introduced by [11] and [10]. A fast alignment-free sequence comparison methods using multiple spaced seeds has been described in [13], see also [24] and [14].

The algorithm, S-conLSH, described in this article is an alignment-free tool designed for mapping of noisy and long reads to the reference genome. The following subsection elaborates the concept of Spaced-context in connection to the proposed algorithm.

## 2 Methods

The algorithm S-conLSH for mapping noisy long reads to the reference genome essentially consists of two steps, reference genome indexing and read mapping. The complete workflow of S-conLSH is provided in Figure 1 and the entire procedure is detailed below.

**Figure 1:**
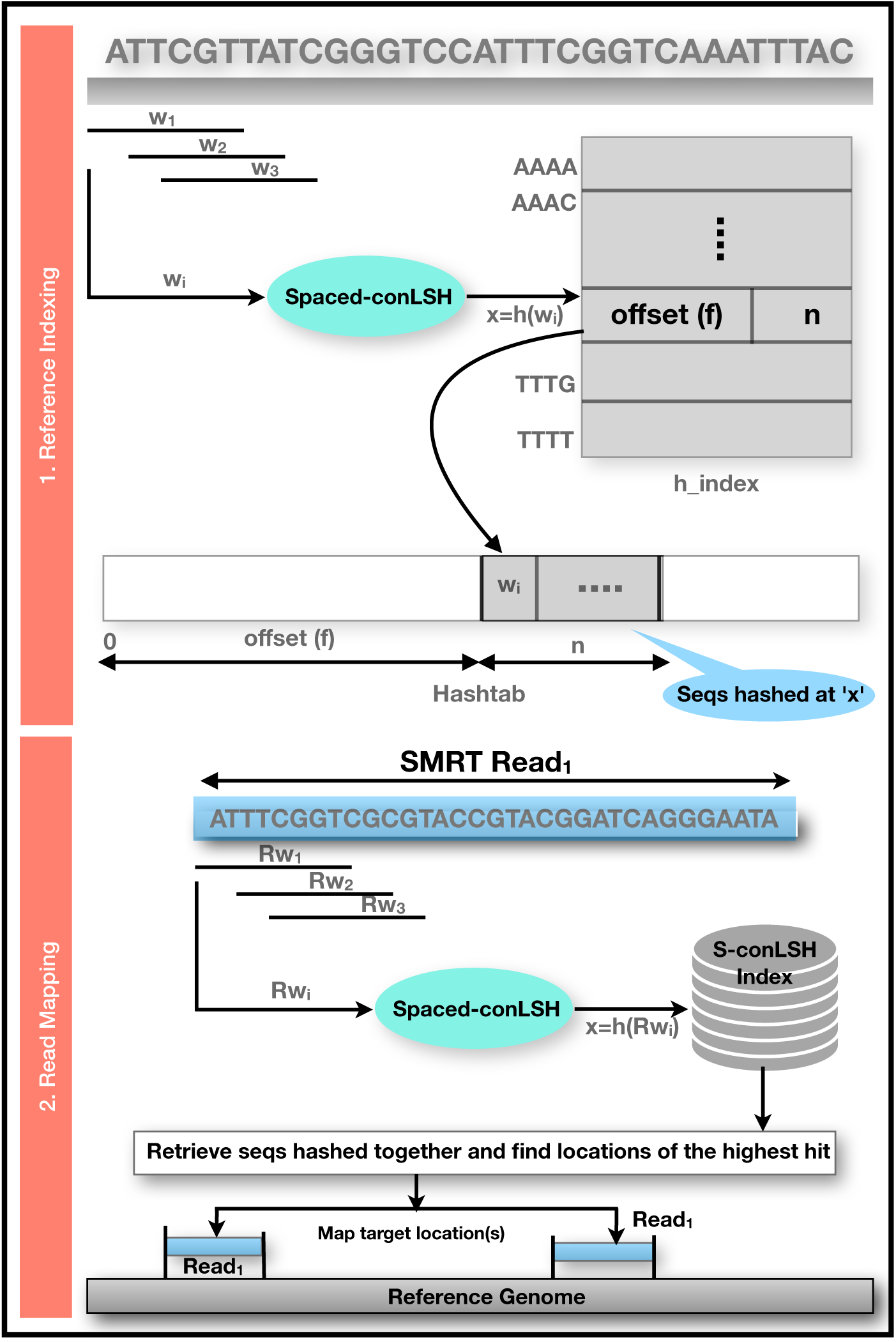
A schematic workflow of Indexing and mapping using S-conLSH.

### 1. Reference Genome Indexing

The reference genome is sliced into overlapping windows, and these windows are hashed into hash tables using suitably designed S-conLSH functions (see Definition 2.5) as shown in Fig. 1. S-conLSH uses two hash tables ‘*h*_*index*’ and ‘*Hashtab*’. An entry in *h*_*index* has two fields (*f, n*): *f* stores an offset to the table *Hashtab*, where sequences are clustered according to their hash values, and *n* is the total number of sequences hashed at a particular value. Therefore, *Hashtab*[*h*_*index*[x]. *f*] to *Hashtab*[*h*_*index*[x]. *f +h*_*index*[x].*n*] are the sequences hashed at value *x*.

### 2. Read Mapping

For each noisy long read, S-conLSH utilizes the same hash function for computing the hash values and retrieves sequences of the reference genome that are hashed in the same position as the read. Finally, the locations of the sequences with the highest hits are chained and reported as an alignment-free mapping of the query read (see Fig. 1).

By default, S-conLSH provides alignment-free mappings of the SMRT reads to the reference genome. If a base level alignment is required, S-conLSH provides an option (--align 1) to generate alignment in SAM format using ksw library (https://github.com/attractivechaos/klib). Some key aspects of S-conLSH are detailed in the following subsections.

### 2.1 Context based Locality Sensitive Hashing

Locality Sensitive Hashing [1, 12] is an approximate near-neighbor search algorithm, where the points having a smaller distance in the feature space, will have a higher probability of making a collision. Under this assumption, a query is compared only to the objects having the same hash value, rather than to all the items in the database. This makes the algorithm work in sublinear time. In the definitions below, we use the following notations:

For a string *x* of length *d* over some set Σ of symbols and 1 ≤ *i* ≤ *j* ≤ *d, x*[*i*] denotes the *i* –th symbol of *x*, and *x*[*i* .. *j*] denotes the (contiguous) substring of *x* from position *i* to position *j*. If *ℋ* is a finite set of functions defined on some set *X*, for any *h* ∈ *ℋ*, randomly drawn with uniform probability, and *x, y* ∈ *X, Pr* _*ℋ*_ [*h*(*x*) *= h*(*y*)] denotes the probability that *h*(*x*) *= h*(*y*).

The definition of Locality Sensitive Hashing as introduced in [1, 12] is given below:

#### Definition 2.1. Locality Sensitive Hashing [1, 12]

Let (*X, D*) be a metric space, let *ℋ* be a family of hash functions mapping *X* to some set *U*, and let *R, c, P*_1_, *P*_2_ be real numbers with *c >* 1 and 0 ≤ *P*_2_ < *P*_1_ ≤ 1. *ℋ* is said to be (*R, cR, P*_1_, *P*_2_)-*sensitive* if for any *x, y* ∈ *X* and *h* ∈ *ℋ*

- *Pr* _*ℋ*_ [*h*(*x*) *= h*(*y*)] ≥ *P*_1_ whenever *D*(*x, y*) ≤ *R*, and
- *Pr* _*ℋ*_ [*h*(*x*) *= h*(*y*)] ≤ *P*_2_ whenever *D*(*x, y*) ≥ *cR*.

To illustrate the concept of locality sensitive hashing for DNA sequences, let us consider a finite set Σ *=* {*A, T,C, G*} called the *alphabet*, together with an integer *d >* 0. Let *X* be the set of all length-*d* words over Σ, endowed with the *Hamming distance*, and let *U* be the alphabet Σ. For 1 ≤ *i* ≤ *d*, let the function *h*_*i*_ : *X* → *U* be defined by *h*_*i*_ (*x*) *= x*[*i*], ∀*x* ∈ *X*. Next, let *R* and *cR* be real numbers with *c >* 1 and 0 ≤ *R* < *cR* ≤ *d*, and define 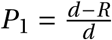 and 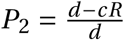. Then the set *ℋ =* {*h*_*i*_ : 1 ≤ *i* ≤ *d*} is (*R, cR, P*_1_, *P*_2_)-sensitive. To see this, observe that for any two words *p, q* ∈ *X*, the probability *Pr* _*ℋ*_ [*h*(*p*) *= h*(*q*)] is same as the fraction of positions *i* with *p*[*i*] *= q*[*i*]. Therefore,

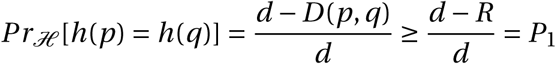

if *D*(*p, q*) ≤ *R*,

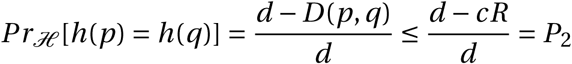

and if *cR* ≤ *D*(*p, q*).

Therefore, *P*_1_ *> P*_2_ as *cR > R*. This proves that the family of hash functions *ℋ =* {*h*_*i*_ : 1 ≤ *i* ≤ *d*} is locality sensitive.

In biological applications, it is often useful to consider the local *context* of sequence positions and to consider matching *subwords*, as shown in conLSH [8]. It groups similar sequences in the localized slots of the hash tables considering the neighborhoods or contexts of the data points. A context in connection to sequence analysis can be formally defined as:

#### Definition 2.2. Context

Let *x* : (*x*_1_*x*_2_ … *x*_*d*_) be a sequence of length *d*. A *context* at the *i*-th position of *x*, for *i* ∈ {*λ +* 1,…, *d* − *λ*}, is a subsequence *x*[*i* − *λ* … *i* … *i + λ*] of length 2*λ+* 1, formed by taking *λ* characters from each of the right and left sides of *x*[*i*]. Here, *λ* is a positive constant termed the *context factor*.

To define context based locality sensitive hashing, the above example is generalized such that, for a given subword length (2*λ* + 1) < *d*, each hash function in *ℋ* will map words containing the same length-(2*λ* + 1) *subwords* at some position to the same bucket in *U*. The subword length (2*λ* + 1) is called the *context size*, where *λ* is the context factor.

#### Definition 2.3. Context based Locality Sensitive Hashing (conLSH)

Let Σ be a set called the *alphabet*. Let *λ* and *d* be integers with (2*λ*+1) < *d*. Let *X* be the set of all length-*d* words over Σ and *U* be the set of all length-(2*λ*+1) words over Σ. For *R, cR, P*_1_, and *P*_2_ as above, a (*R, cR, P*_1_, *P*_2_)-*sensitive* family *ℋ* of functions mapping *X* to *U* is called (*R, cR, λ, P*_1_, *P*_2_)-*sensitive*, if for each *h* ∈ *ℋ*, there are positions *i*_*h*_ and *j*_*h*_ with *λ*+ 1 ≤ *i*_*h*_, *j*_*h*_ ≤ *d* − *λ* such that for all *p, q* ∈ *X* one has *h*(*p*) *= h*(*q*) whenever

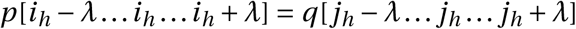

holds.

### 2.2 Gapped read Mapping using Spaced-context based Locality Sensitive Hashing

The proposed method S-conLSH, uses spaced-seeds or patterns of 0’s and 1’s in connection with S-conLSH function. For a pattern *𝒫*, the *spaced-context* of a DNA sequence can be defined as:

#### Definition 2.4. Spaced-context

Let *𝒫* be a binary string or *pattern* of length *ℓ*, where ‘1’ represents *match position* and ‘0’ represents *don’t-care position*. Let *ℓ*_*w*_ denote the weight of *𝒫* which is equal to the number of ‘1’s in the pattern. Evidently, *ℓ*_*w*_ ≤ *ℓ*. Let *x* be a sequence of length *d* over alphabet {*A, T,G,C*} such that *ℓ* ≤ *d*. Then, a string *sw* over {*A, T,G,C*} of length *ℓ*_*w*_ is called a *spaced-context of x with respect to 𝒫*, if *sw* [*i*] *= x*[*j*] holds if and only if *𝒫* [*j*] *=* 1, where *i* ≤ *j*, 1 ≤ *i* ≤ *ℓ*_*w*_ and 1 ≤ *j* ≤ *ℓ*.

Sequences sharing a similar spaced-context with respect to a pre-defined pattern *P*, are hashed together in S-conLSH.

The concept of gap-amplification is used in locality sensitive hashing to ensure that the dissimilar items are well separated from the similar ones. To do this, gap between the probability values *P*_1_ and *P*_2_ needs to be increased. This is achieved by choosing *L* different hash functions, *g*_1_, *g*_2_, …, *g*_*L*_, such that *g* _*j*_ is the concatenation of *K* randomly chosen hash functions from *ℋ*, i.e., *g* _*j*_ *=* (*h*_1,*j*_, *h*_2,*j*_, …, *h*_*K, j*_), for 1 ≤ *j* ≤ *L*. This procedure is known as “gap amplification” and *K* is called the “concatenation factor” [1]. For every hash function *g* _*j*_, 1 ≤ *j* ≤ *L*, there is a pattern *𝒫*_*j*_ associated with it. The *spaced-context based Locality Sensitive Hashing* is now defined as follows:

#### Definition 2.5. Spaced-context based Locality Sensitive Hashing (S-conLSH)

Let *sw*_*j*_ (*x*) be the spaced-context of sequence *x* with respect to the binary pattern *P*_*j*_ of length *ℓ*, 1 ≤ *j* ≤ *L*. Let *𝒫*_*j*_ be defined by the regular expression (0^*^(1)^(2*λ*+1))^ *K* 0^*^. Therefore, the weight of *𝒫*_*j*_, i.e., *ℓ*_*w*_ *=* (2*λ*+ 1)*K*. The maximum value of *ℓ* would be (2*λ*+ 1)*K + z*(*K*+ 1) assuming that at most *z* zeros are present between two successive contexts of 1’s in *𝒫*_*j*_, where *z* ≥ 0 is an integer parameter. Let *d* be an integer with *ℓ* ≤ *d, X* be the set of all length-*d* words over Σ and *U* be the set of all length-*ℓ*_*w*_ words over Σ. For *R, cR, P*_1_, *P*_2_, and *λ* as introduced in Definition 2.3, a (*R, cR, λ, P*_1_, *P*_2_)-*sensitive* hash function *g* _*j*_ *=* (*h*_1,*j*_, *h*_2,*j*_, …, *h*_*K, j*_), where *h*_*i, j*_ ∈ *ℋ*, 1 ≤ *i* ≤ *K*, mapping *X* to *U* is called (*R, cR, λ, z, P*_1_, *P*_2_)-*sensitive*, if for any *p, q* ∈ *X* one has *g* _*j*_ (*p*) *= g* _*j*_ (*q*) whenever *sw*_*j*_ (*p*) *= sw*_*j*_ (*q*) holds with respect to the pattern *𝒫*_*j*_.

#### Algorithm 1 Spaced-conLSH pattern Generation(*L,K, λ,z*)

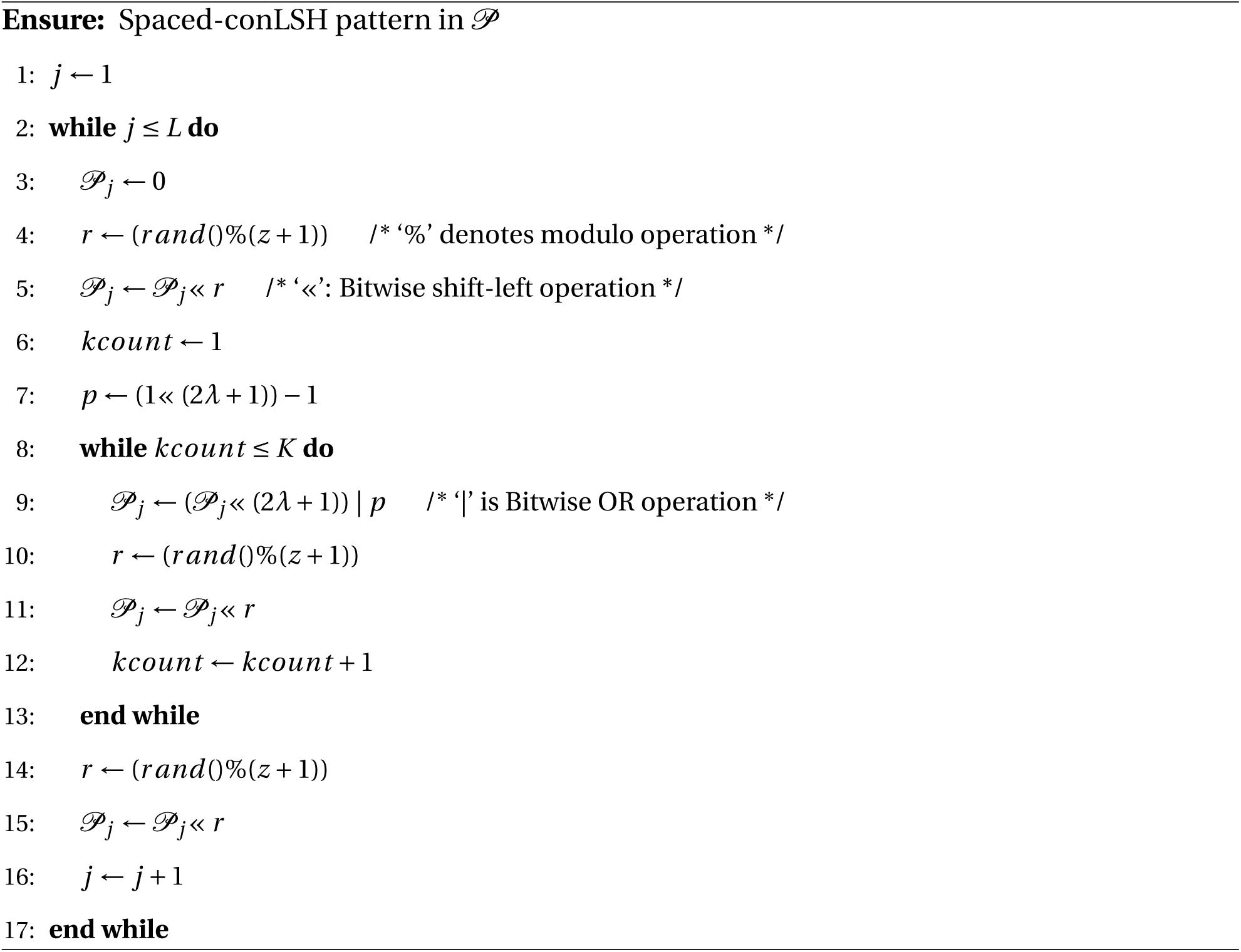

Therefore, instead of restricting similarity over the (2*λ* + 1)*K* consecutive bases as was done for conLSH [8], S-conLSH incorporates greater flexibility by checking only the positions which correspond to a 1 in the pattern. For example, the binary string “011100111” is a pattern for a system having *K =* 2, *z =* 2 and context size (2*λ*+ 1) *=* 3. The hash value or the spaced-context of the string “ATTCGGTAA” for the above pattern will be “TTCTAA” (see Figure 2(b)). In S-conLSH, noisy long reads are hashed using *L* functions corresponding to *L* different patterns generated using Algorithm 1. Multiple pattern based functions enable gapped-mapping of the reads as illustrated in Figure 2. Consider a scenario of two patterns *𝒫*_1_ *=*“011100111” and *𝒫*_2_ *=*“111111” having context size *=* 3, *L =* 2 and *K =* 2. The string *p =*“ATTCGGTAA” generates two hash values *sw*_1_(*p*) *=*“TTCTAA” and *sw*_2_(*p*) *=*“ATTCGG” for the patterns *𝒫*_1_ and *𝒫*_2_ respectively (see Figure 2(b)). Similarly, *sw*_1_(*q*) *=*“TCTGTA” and *sw*_2_(*q*) *=*“TTCTAA” are the hash values for string *q =*“TTCTAAGTA” (Figure 2(c)). As shown in the hash table of Figure 2(d), the two strings collide to the same bucket of the hash table due to the common hash value “TTCTAA”. This results in mapping with three gaps or indels, corresponding to the three 0’s of “011100111”, in the second string. This gapped-mapping is a powerful feature of S-conLSH which is quite uncommon in standard spaced-seed based methods.

**Figure 2:**
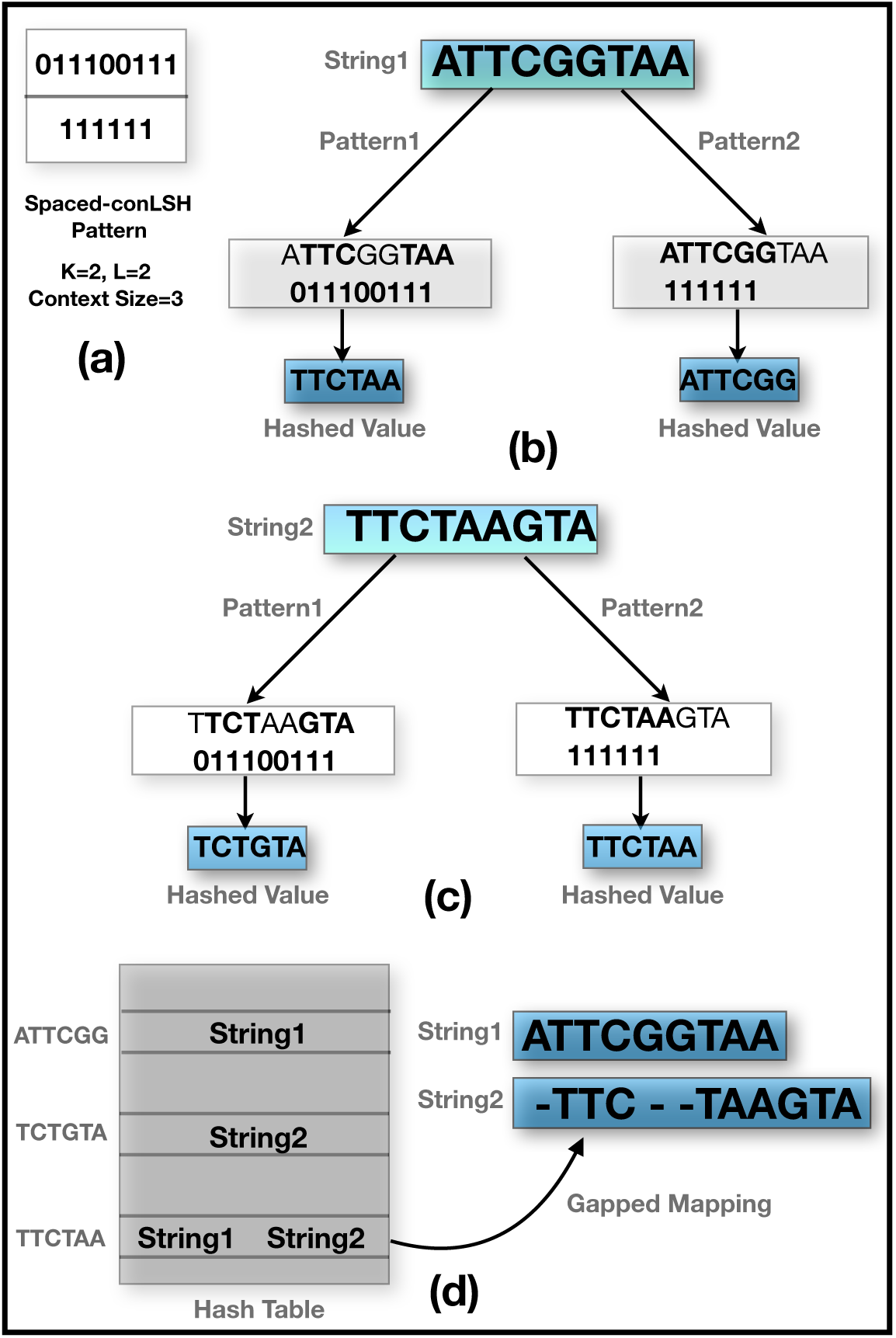
A schematic illustration of gapped-mapping using S-conLSH. (a) Multiple patterns having context size *=* 3 and *K =* 2. (b) & (c) Hashing of the strings “ATTCGGTAA” and “TTCTAAGTA” respectively using different patterns. (d) Final hash table and gapped-mapping of the two strings due to the collision at “TTCTAA”

To obtain an integer hash value from the Spaced-context, an encoding function 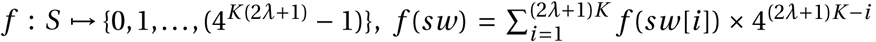, ∀*sw* ∈ *S*, has been defined assuming *f* (*A*) *=* 0, *f* (*C*) *=* 1, *f* (*G*) *=* 2 and *f* (*T*) *=* 3, where *S* is the set of all spaced-contexts of length (2*λ*+ 1)*K* defined over the alphabet {*A, T,C, G*}. A pattern produces hash values of length equal to its weight. Keeping the weight same, the pattern length is increased in S-conLSH by introducing don’t care positions (or, zeros). This allows S-conLSH to look at a larger portion of the sequences without increasing the computational overhead. Consequently, conLSH is able to find distant homologs that might otherwise be overlooked. Not only that, it provides better sensitivity in resolving repeats because of the consideration of the neighborhood (or, contexts) when measuring the similarity between the sequences. S-conLSH has a provision of split mapping for chimeric reads as follows. If a read fails to get associated with end-to-end mapping, it is split into a series of non-overlapping segments and re-hashed to find target location(s) for each segment.

## 3 Results

Six different real and simulated datasets of *E.coli, A.thaliana, O.sativa, S.cerevisiae* and *H.sapiens* have been used to benchmark the performance of S-conLSH in comparison to other state-of-the art aligners, *viz*., Minimap2 [17], lordFAST [9], Minimap [16], conLSH [8] and MUMmer4 [23]. All these methods are executed in a setting designed for PacBio read alignment (see Table 1) in single-threaded mode. The default parameter settings used for S-conLSH are *K =* 2, context size (2*λ*+ 1) *=* 7, *L =* 2 and *z =* 5. The datasets used in the experiment have been summarized in Table 2.

**Table 1:** Parameter settings and commands used by different methods for mapping of PacBio SMRT reads.

| Mapper | Command line settings |
| --- | --- |
| Minimap [16] | `/minimap -d in.mmi ref_file ./minimap -l in.mmi read_file > output_file` |
| Minimap2 [17] | `./minimap2 -Hk19 -d in.mmi ref_file ./minimap2 -a -Y -x map-pb -t 1 in.mmi read_file > output_file` |
| lordFAST [9] | `./lordfast --index ref_file ./lordfast --search ref_file --seq read_file --thread 1 > output_file` |
| MUMmer4 [23] | `./nucmer --sam-long=output_file ref_file read_file -l 15 -c 31` |
| conLSH [8] | `./conLSH-indexer $PATH ref_file -K 2 -L 1 > output_file ./conLSH-aligner $PATH read_file ref_file -K 2 -L 1 -l 8 -w 1000 -m 5 --lambda 2 > output_file` |
| S-conLSH | `./S-conLSH $PATH ref_file read_file -K 2 --lambda 3 -L 2 -z 5 > output_file` |

**Table 2:** The summary of real and simulated datasets used in the experiment along with the corresponding reference genome links

| Dataset | Type | Platform | # of reads | Reference Genome |
| --- | --- | --- | --- | --- |
| H. sapiens-real | real | PacBio RS II P5/C3 release | 290992 | hg38 |
| E. coli-real | real | PacBio RS II P5/C3 release | 300 | Escherichia coli str. K-12 substr. MG1655 |
| A. thaliana-real | real | PacBio RS II P5/C3 release | 3448228 | TAIR10 |
| O.sativa-real | real | PacBio RS II P5/C3 release | 590268 | Build 4.0 |
| S.cerevisiae-real | real | PacBio RS II P5/C3 release | 594243 | S288C (assembly R64) |
| H. sapiens-sim | simulated | PBSIM | 146932 | hg38 |

The aligner, rHAT [20] has been excluded from the study, as it has been reported to malfunction in certain scenarios [17]. The PacBio read alignment module of BWA-MEM [15] has been replaced by Minimap2, as it retains all the main features of BWA-MEM, while being 50×faster and more accurate. Therefore, the results of BWA-MEM are not shown separately in the tables. Moreover, BLASR [6] has also not been used in the comparative study, as Minimap2 and lord-FAST have been found to outperform it in all respect.

By default, S-conLSH produces output in pairwise read mapping format (PAF) ([16]). There are scripts available to convert the popular SAM [18] alignment formats to PAF ([16]). If a base-to-base alignment is requested, S-conLSH provides an option (--align 1), where the target locations are aligned using ksw alignment library (https://github.com/attractivechaos/klib) to produce the SAM file. The entire experiment has been conducted on an Intel Core i7-6200U CPU @ 2.30GHz × 16(cores), 64-bit machine with 32GB RAM.

The results demonstrated in this article are organized into three categories: i) Performance on simulated datasets, ii) Study on real PacBio reads, and iii) Robustness of S-conLSH for different parameter settings.

### 3.1 Experiment on Simulated Dataset

To study the accuracy of SMRT read mapping, a total of 146932 noisy long reads have been simulated from hg38 human genome using PBSIM [25] command “`pbsim --data-type CLR --depth 1 --length-min 1 --length-max 200000 --seed 0 --sample-fastq real.fastq hg38.fa`”. The error profile has been sampled from three real human PacBio RS II P5/C3 reads listed below, concatenated as `real.fastq`.

- `m130929_024849_42213_c100518541*_s1_p0.1.subreads.fastq`
- `m130929_024849_42213_c100518541*_s1_p0.2.subreads.fastq`
- `m130929_024849_42213_c100518541*_s1_p0.3.subreads.fastq`

The simulated reads from 5 different Human chromosomes are used to test the performance of S-conLSH in comparison to the other standard aligners. The sensitivity and precision have been computed based on the ground truth as obtained from the .maf files of PBSIM. A read is considered to be mapped correctly (as defined by [9]) if i) it gets mapped to the correct chromosome and strand; and ii) the target subsequence of reference genome where the read maps to, must overlap with the true mapping by at least 1bp. The sensitivity is measured as a fraction of correctly mapped reads out of the total number of reads. Precision is defined, in the same way, as the fraction of correctly mapped reads out of the total number of mapped reads.

Table 3 summarizes the number of correct mappings, sensitivity, precision, and running time by different methods, Minimap, Minimap2, lordFAST, MUMmer4, conLSH, and S-conLSH for a total of 146932 reads simulated from five different human chromosomes. The number of reads extracted from each chromosome is listed in Table 3. The result shows that S-conLSH produces the highest number of correct mappings among all five aligners for different chromosomes of Human-sim dataset. S-conLSH maps 32111 reads out of total 32290 reads of Chr#1, among which 31964 mappings are found to be correct when compared with the ground truth. Minimap2 is the second highest in producing the correct mappings in this case. A similar scenario has been generally observed for the four other chromosomes as well. It is clear that Minimap2 always aligns all the reads to some location in the reference genome, but produces more incorrect mappings when compared to S-conLSH. Evidently, S-conLSH provides the highest sensitivity for all the chromosomes considered. Minimap, on the other hand, exhibits higher precision but lower sensitivity as it leaves a large number of reads unaligned. The number of unaligned reads by Minimap increases for large and complicated real datasets (see Subsection 3.2).

As can be seen, S-conLSH takes 38 CPU seconds to map the reads of chromosome 1, which is slightly slower than Minimap. The speed of Minimap is achieved as it maps a smaller number of reads compared to other aligners. Interestingly, S-conLSH has been found to have smaller mapping time than all the remaining algorithms, while having the maximum number of correctly mapped reads. As there was no separate indexing and aligning time available for MUMmer4, the total time is mentioned as “Mapping time”. MUMmer4 has been found to consume a large amount of time to achieve a desired level of sensitivity. It is evident from Table 3 that the indexing time is quite low for both Minimap and Minimap2. Indexing time for S-conLSH is relatively higher, though it is much smaller as compared to lordFAST. Here, it may be noted that indexing is performed only once for a given reference genome, while the read mapping will need to be performed every time a different individual is sequenced. The compressed and memory-efficient B-tree indexing of conLSH makes it the fastest in processing of reference genomes. However, the mapping time of conLSH is large as it performs base-to-base alignments using Sparse Dynamic Programming. The stringent ungapped matching requirement of the aligner over the entire context of the sequences results in lower sensitivity, after lordFAST. The proposed alignment-free tool, S-conLSH, has been found to be useful in such cases as it obtains the gapped mapping of the noisy reads using spaced-contexts.

**Table 3:** Comparative study of the number of correct mappings, sensitivity, precision, and running time by different methods, Minimap, Minimap2, lordFAST, MUMmer4, conLSH, and S-conLSH, for a total of 146932 reads simulated from five different human chromosomes.

| Chr# | #Reads | Mapper | #Mapped Reads | #Correct Mapping | Sensitivity(%) | Precision(%) | Indexing Time (sec) | Mapping Time (sec) |
| --- | --- | --- | --- | --- | --- | --- | --- | --- |
| Chr1 | 32290 | Minimap | 31591 | 31585 | 97.82 | **99.99** | 15 | 30 |
|  |  | Minimap2 | **32290** | 31863 | 98.69 | 98.69 | 10 | 61 |
|  |  | lordFAST | **32290** | 29313 | 90.79 | 90.79 | 192 | 206 |
|  |  | MUMmer4 | 31940 | 31645 | 98.01 | 99.08 | - | 310 |
|  |  | conLSH | 31945 | 29620 | 91.73 | 92.72 | 08 | 235 |
|  |  | S-conLSH | 32111 | **31964** | **99** | 99.55 | 51 | 38 |
| Chr2 | 34309 | Minimap | 33623 | 33613 | 97.98 | **99.98** | 16 | 33 |
|  |  | Minimap2 | **34309** | 33864 | 98.71 | 98.71 | 10 | 64 |
|  |  | lordFAST | **34309** | 31173 | 90.87 | 90.87 | 170 | 216 |
|  |  | MUMmer4 | 34056 | 33914 | 98.85 | 99.59 | - | 312 |
|  |  | conLSH | 34082 | 31230 | 91.03 | 91.63 | 10 | 229 |
|  |  | S-conLSH | 34153 | **34008** | **99.13** | 99.58 | 54 | 44 |
| Chr3 | 28109 | Minimap | 27481 | 27477 | 97.76 | **99.99** | 15 | 25 |
|  |  | Minimap2 | **28109** | 27698 | 98.55 | 98.55 | 8 | 52 |
|  |  | lordFAST | **28109** | 25513 | 90.77 | 90.77 | 135 | 167 |
|  |  | MUMmer4 | 27894 | 27791 | 98.87 | 99.64 | - | 253 |
|  |  | conLSH | 27899 | 25603 | 91.08 | 91.77 | 07 | 198 |
|  |  | S-conLSH | 27957 | **27863** | **99.13** | 99.67 | 45 | 30 |
| Chr4 | 26871 | Minimap | 26307 | 26301 | 97.88 | **99.98** | 16 | 23 |
|  |  | Minimap2 | **26871** | 26501 | 98.63 | 98.63 | 8 | 51 |
|  |  | lordFAST | **26871** | 24403 | 90.82 | 90.82 | 129 | 158 |
|  |  | MUMmer4 | 26638 | 26533 | 98.75 | 99.61 | - | 248 |
|  |  | conLSH | 26650 | 24449 | 90.98 | 91.74 | 07 | 180 |
|  |  | S-conLSH | 26748 | **26650** | **99.18** | 99.64 | 41 | 29 |
| Chr5 | 25353 | Minimap | 24859 | 24849 | 98.02 | **99.96** | 14 | 21 |
|  |  | Minimap2 | **25353** | 25056 | 98.84 | 98.84 | 7 | 48 |
|  |  | lordFAST | **25353** | 23069 | 91 | 91 | 123 | 149 |
|  |  | MUMmer4 | 25126 | 24951 | 98.42 | 99.31 | - | 234 |
|  |  | conLSH | 25149 | 23106 | 91.14 | 91.87 | 06 | 167 |
|  |  | S-conLSH | 25242 | **25155** | **99.22** | 99.66 | 39 | 23 |

### 3.2 Experiment on Real PacBio Datasets

This section demonstrates the performance of S-conLSH in comparison to other state-of-the-art aligners on five different SMRT datasets of *E.coli-real, A.thaliana-real, O.sativa-real, S.cerevisiae-real* and *H.sapiens-real* (refer Table 2 for details). A comparative study of running time, percentage of reads aligned and coverage by different aligners for real human SMRT subread named `m130929_024849_42213_c100518541*_s1_p0.1.subreads .fastq` consisting of 23235 reads has been included in Table 4. Results on MUMmer4 are excluded since it takes inordinately long.

**Table 4:** Comparative study of running time, percentage of reads aligned and coverage by different aligners for *H.sapiens-real* SMRT dataset of 23235 reads.

| Mapper | Indexing Time (Sec) | Mapping Time (Sec) | % of Reads Aligned | Mean Coverage |
| --- | --- | --- | --- | --- |
| Minimap | 140 | 30 | 94.8 | NA |
| Minimap2 | 138 | 106 | 100 | 0.0473 |
| lordFAST | 2286 | 327 | 100 | 0.0566 |
| conLSH | 47 | 404 | 99 | 0.0579 |
| S-conLSH | 794 | 99 | 99.9 | 0.078 |

It can be seen that S-conLSH provides the highest coverage value among the five standard methods used in the experiment. Minimap does not have any coverage statistics, as it is unable to produce alignment as SAM file. The performance in terms of indexing and mapping time, as shown in Table 4, is similar to that has already been observed for simulated datasets. The percentage of read alignment is the highest by Minimap2 and lordFAST. This is similar to the scenario obtained on simulated datasets where Minimap2 and lordFAST align all the reads against the reference genome, even though it may contain some incorrect mappings. S-conLSH, on the other hand, has a mapping ratio of 99.9%, which is lower than Minimap2 and lordFAST. This is due to the fact that S-conLSH gives higher priority to the mapping accuracy and it leaves a few reads unaligned if potential target locations are not found. S-conLSH has a higher memory footprint of about 13GB for indexing the entire human genome.

Similar results are observed for *E.coli-real, A.thaliana-real, O.sativa-real*, and *S.cerevisiaereal* real PacBio datasets as can be seen in Table 5. It is clear that S-conLSH is among the fastest in terms of mapping time, after Minimap. However, Minimap fails to align a good portion of the reads for large datasets like *A.thaliana* and *S.cerevisiae*. The aligner, conLSH, on the other hand, requires lower indexing time but higher mapping time to align a reasonable amount of reads to the reference genome. However, the alignment quality of conLSH is often compromised as studied in Section 3.1. The percentage of reads mapped by S-conLSH is generally lower than Minimap2 and lordFAST, as it tries to ensure the best of the mapping quality. It is, however, difficult to measure the quality of read mappings for real datasets, as there is no ground truth available for such cases.

**Table 5:** Comparative study of running time, percentage of reads aligned by different aligners for four datasets of *E.coli-real, A.thaliana-real, O.sativa-real* and *S.cerevisiae-real*.

| dataset | #Reads | Mapper | Index Time (Sec) | Mapping Time (Sec) | % of Reads Aligned |
| --- | --- | --- | --- | --- | --- |
| *E.coli-real* | 300 | Minimap | 1 | 1 | 91.6 |
|  |  | Minimap2 | 2 | 2 | 100 |
|  |  | lordFAST | 3 | 10 | 100 |
|  |  | conLSH | 1 | 2 | 100 |
|  |  | S-conLSH | 2 | 1 | 98.3 |
| *A.thaliana-real* | 3448228 | Minimap | 7 | 798 | 88 |
|  |  | Minimap2 | 6 | 10562 | 100 |
|  |  | lordFAST | 78 | 24277 | 100 |
|  |  | conLSH | 1 | 40028 | 99.4 |
|  |  | S-conLSH | 27 | 2073 | 93.7 |
| *O.sativa-real* | 590268 | Minimap | 20 | 487 | 94.3 |
|  |  | Minimap2 | 20 | 6898 | 100 |
|  |  | lordFAST | 287 | 10462 | 100 |
|  |  | conLSH | 6 | 18024 | 97 |
|  |  | S-conLSH | 88 | 962 | 98.2 |
| *S.cerevisiae-real* | 594243 | Minimap | 1 | 104 | 41.8 |
|  |  | Minimap2 | 1 | 881 | 100 |
|  |  | lordFAST | 7 | 2130 | 100 |
|  |  | conLSH | 1 | 7890 | 99.8 |
|  |  | S-conLSH | 3 | 299 | 90.3 |

### 3.3 Robustness of S-conLSH for Different Parameter Settings

An exhaustive experiment with different values of *K, λ, L*, and *z* has been carried out to study the robustness of the proposed method S-conLSH.

Table 6 summarizes the study of indexing and mapping time along with the percentage of reads aligned for different values of S-conLSH parameters on real human SMRT dataset `m130929_024849_42213_c100518541*_s1_p0.1.subreads.fastq`. As can be observed the best performance (highest percentage of read mapping in minimum time) is achieved with the settings *K =* 2, (2*λ*+ 1) *=* 7, *L =* 2 and *z =* 5. The mapping time increases with *L* as it directly corresponds to the number of hash tables used to retrieve the target locations. Indexing time, on the other hand, is proportional to the product (2*λ*+ 1)*K*. It seems that *z* has little effect on the performance of S-conLSH as the running time and the percentage of reads aligned mostly stay invariant with *z*. However, the parameter *z* is important to enhance the sensitivity of the method. This is reflected in Table 7 when studied on simulated reads. The highest number of correct mappings is obtained with the default settings (shown in boldface) when *z =* 5. The zeros in the spaced-seed help to find the distant similarities as it encompasses a larger portion of the sequence while the weight ((2*λ*+ 1)*K*) of the pattern remains the same. However, a very large value of *z* may degrade the accuracy as it joins unrelated contexts together. It is evident from Tables 6 and 7 that the performance of S-conLSH remains reasonably good irrespective of the variation of the parameter values. Therefore, it can be concluded that the algorithm SconLSH is quite robust even though it requires some tuning of different parameters for the best performance.

**Table 6:** Performance of conLSH with change of *K, L, z*, and *λ* for real human SMRT dataset. The default setting is marked as bold.

| Concatenation<br>Factor ($K$) | Context Size<br>$2 \times \lambda + 1$ | $L$ | $z$ | Indexing<br>Time(s) | Mapping<br>Time(s) | % of Reads<br>Mapped |
| --- | --- | --- | --- | --- | --- | --- |
| 2 | 7 | 1 | 5 | 790 | 80 | 97.7 |
| **2** | **7** | **2** | **5** | **794** | **99** | **99.9** |
| 2 | 7 | 3 | 5 | 797 | 122 | 99.9 |
| 2 | 7 | 1 | 7 | 791 | 82 | 97.7 |
| 2 | 7 | 1 | 11 | 794 | 80 | 97.7 |
| 2 | 5 | 1 | 5 | 67 | 11 | 73.4 |
| 4 | 3 | 1 | 5 | 302 | 107 | 97.2 |
| 3 | 5 | 1 | 5 | 849 | 85 | 96.8 |

**Table 7:**
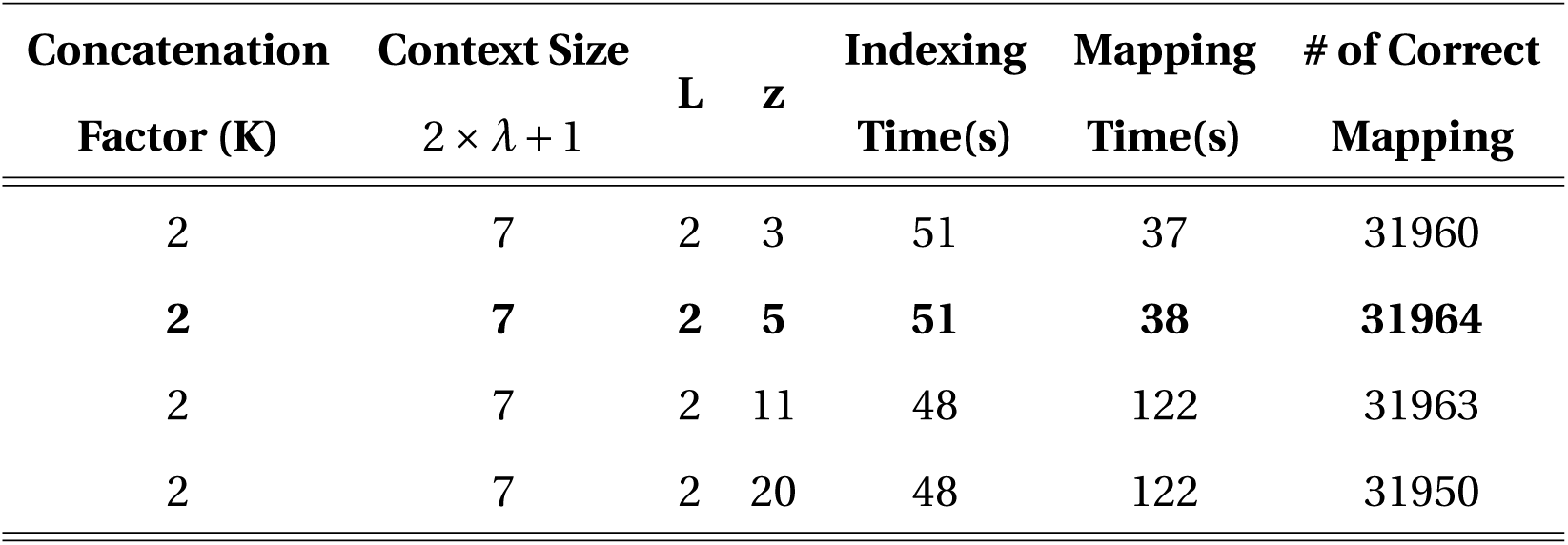
Performance of conLSH with change of *z* for *chromosome 1* of *H.sapiens-sim* dataset consisting of 32290 reads. The default parameter setting is shown in boldface.

## 4 Discussion and Conclusions

S-conLSH is one of the first alignment-free reference genome mapping tools achieving a high level of sensitivity. Earlier, Minimap was designed to map reads against the reference genome without performing an actual base-to-base alignment. However, the low sensitivity of Minimap precluded its applications in real-life domains. Minimap2 is one of the best performing state-of-the-art alignment-based methods which provides an excellent balance of running time and sensitivity. The method described in this article, S-conLSH, has been observed to outperform Minimap2 in respect of sensitivity, precision, and mapping time. However, it has a longer indexing time and a higher memory footprint. Nevertheless, sequence indexing is a one-time affair, and memory is inexpensive nowadays.

The *spaced*-context in S-conLSH is especially suitable for extracting distant similarities. The variable-length spaced-seeds or patterns add flexibility to the proposed algorithm. Multiple patterns (with higher values of *L*) increase the sensitivity but at the cost of more time. Moreover, with the introduction of don’t care positions, the patterns become longer, thus providing better performance in resolving conflicts that occur due to the repetitive regions. The provision of rehashing for chimeric read alignment and reverse strand mapping make S-conLSH ideal for applications in the real-life sequence analysis pipeline.

A memory-efficient version of the S-conLSH can be developed in the future. The algorithm, at its current stage, can not conclude on the optimal selection of the patterns. A study on finding the optimal set of spaced-seeds can be carried out in future to improve the performance of the algorithm. Though the experiment demonstrated in this article is confined to the noisy long reads of PacBio datasets, it can be further extended on ONT reads as well. Finally, we would like to conclude with a strong expectation that the proposed method S-conLSH will draw the attention of the peers as one of the best performing reference mapping tools designed so far.

